# Proteomics and single-cell transcriptomics identify LRP1 as a diagnostic biomarker for atypical fibroxanthoma and pleomorphic dermal sarcoma

**DOI:** 10.64898/2026.02.04.703892

**Authors:** Joshua J. Lingo, Maggie M. Balas, Arjun M. Bashyam, Gregory A. Hosler, Galen T. Squiers, Jason C. Klein

## Abstract

Atypical fibroxanthoma (AFX) and pleomorphic dermal sarcoma (PDS) are cutaneous neoplasms that fall along a spectrum. PDS is more aggressive than AFX with higher rates of local and distant metastases. Diagnostic biomarkers for AFX and PDS are lacking and therefore these tumors are diagnosed only after excluding other dermal spindle cell neoplasms, including cutaneous leiomyosarcoma (cLMS), spindle cell melanoma (SCM), and sarcomatoid squamous cell carcinoma (sSCC). To identify clinically valuable biomarkers, we contrast the tumors within the diagnostic differential using single-cell RNA sequencing and bulk proteomic data. Gene Ontology (GO) analysis of transcripts and proteins enriched in AFX/PDS identified multiple shared pathways associated with cell adherence and the extracellular matrix. We identify that LRP1, LTBP2, and NAV1 are all enriched in AFX/PDS over other tumors in the differential at both the level of mRNA and protein. IHC reveals that LRP1 is 90% sensitive and 73% specific for AFX/PDS in a cohort of AFX, PDS, cLMS, SCM, and sSCC. This outperforms published data for CD10, which is currently used clinically (sensitivity 83.5% and specificity 50%). When used in conjunction with LTBP2, specificity for AFX/PDS within the differential rises from 73% to 93%. These findings suggest that LRP1, particularly if evaluated in conjunction with existing stains, can improve diagnostic accuracy for AFX and PDS.

## To the editor

Atypical fibroxanthoma (AFX) and pleomorphic dermal sarcoma (PDS) are two cutaneous tumors that fall along a spectrum, with AFX being confined to the dermis and indolent while PDS invades into the subcutis and has higher rates of local recurrence and metastasis (1, 2). Currently, AFX/PDS is diagnosed after excluding other atypical dermal spindle cell tumors including sarcomatoid squamous cell carcinoma (sSCC), cutaneous leiomyosarcoma (cLMS), and desmoplastic/spindle cell melanoma (SCM). CD10 and procollagen-1 are sensitive biomarkers for AFX and PDS (70-90% sensitivity); however, specificity is limited by positive staining in SCM, sSCC, and cLMS (30-60% specificity) (3-5). Positive biomarkers must be interpreted in conjunction with additional immunohistochemical stains to rule out other dermal spindle cell neoplasms. To better understand these tumors, we recently performed the first single-cell RNA sequencing (scRNA-seq) of AFX and PDS, identifying *COL6A3* as a prognostic biomarker (6, 7).

Here, we compare the transcriptional profiles of AFX and PDS against published scRNA-seq from other tumors in the AFX/PDS differential (Figure 1A and 1B) (6, 8-10). We perform differential gene expression analysis to identif y the top 100 upregulated genes in AFX/PDS tumor cells compared to cutaneous squamous cell carcinoma (cSCC), melanoma, and leiomyosarcoma. We highlight multiple biological processes (Figure 1C) and molecular functions (Figure 1D) related to cell adherence, collagen, and extracellular matrix remodeling through gene ontology (GO) analysis of these 100 genes. To narrow the pool of candidate biomarkers, we filtered our list for transcripts that are upregulated in AFX/PDS relative to healthy skin (11). After filtering, 12 candidate biomarkers remained: *LUM, FBN1, COL5A2, COL5A3, CDH11, BMP1, LRP1, LOXL2, NAV1, DPYSL3, PHLDB1*, and *LTBP2* (Supplemental Figure 1).

**Figure 1:**
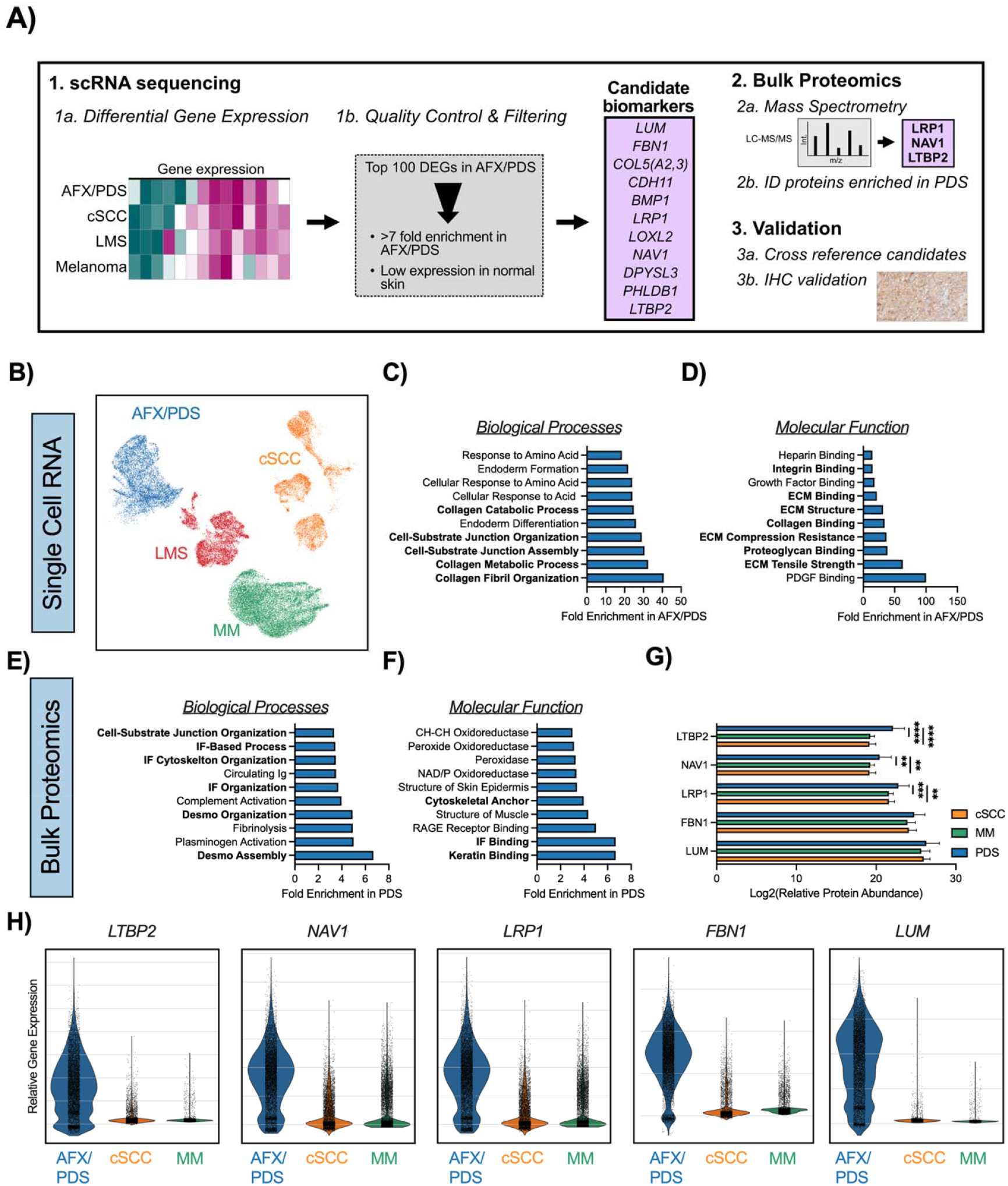
Single-cell RNA sequencing and bulk proteomic analyses of dermal spindle cell neoplasms. A) Schematic of bioinformatic workflow to analyze single-cell RNA sequencing (scRNA-seq) and proteomic data with immunohistochemical validation. B) Dimensional reduction and visualization of scRNA-seq with AFX/PDS in blue, cSCC in orange, LMS in red, and MM in green. Gene Ontology (GO) analysis of the top 100 upregulated genes in AFX/PDS revealed enriched C) biological processes and D) molecular functions. Gene Ontology (GO) analysis of upregulated proteins in AFX/PDS revealed enriched E) biological processes and F) molecular functions. G) Relaive protein abundance of select candidate biomarkers in PDS, MM, and cSCC. H) Violin plots illustrating relative gene expression for select candidate biomarkers in AFX/PDS, cSCC, and MM.

To investigate the utility of these transcriptional targets as specific immunohistochemistry markers in the AFX/PDS differential, we cross-reference candidates against bulk proteomic data including PDS, cSCC, and malignant melanoma (12). We identify that proteins upregulated in PDS are enriched for biological processes (Figure 1E) and molecular functions (Figure 1F) related to cell-cell and cell-junction adherence, similar to the GO analysis of upregulated transcripts. Of the 12 candidate biomarkers, only five were included in the proteomic dataset. *LRP1, LTBP2*, and *NAV1* have significantly increased protein expression compared to cSCC and malignant melanoma (Figure 1G). *FBN1* and *LUM* are modestly upregulated in PDS at the protein level but failed to reach statistical significance and were excluded from downstream analysis. We highlight relative gene expression of all five candidates in Figure 1H.

To determine if our candidates can be used as diagnostic biomarkers for AFX/PDS, we established a cohort of dermal spindle cell neoplasms for immunohistochemical validation including 5 AFX, 5 PDS, 5 cLMS, 5 SCM, and 5 sSCC (Figure 2A). Immunohistochemistry reveals that LRP1 is positive in 90% of AFX/PDS, 60% of cLMS, 0% of SCM, and 20% of SSCC (Figure 2B), with an overall sensitivity of 90% and specificity of 73%. Reciever Operating Characteristic (ROC) curves indicate that LRP1 can accurately diagnose AFX and PDS (AUC: AFX/PDS vs differential = 0.817) (Figure 2C) and outperforms the most common clinically used stain, CD10 (ROC AUC 0.817 vs 0.667) (Figure 2D) (5). Additionally, LTBP2 is positive in 60% of AFX/PDS, 20% of cLMS, 20% of SCM, and 20% of sSCC (Supplemental Fig 2A-B), with an overall sensitivity of 60%, specificity of 80%, and ROC AUC of 0.7.

**Figure 2:**
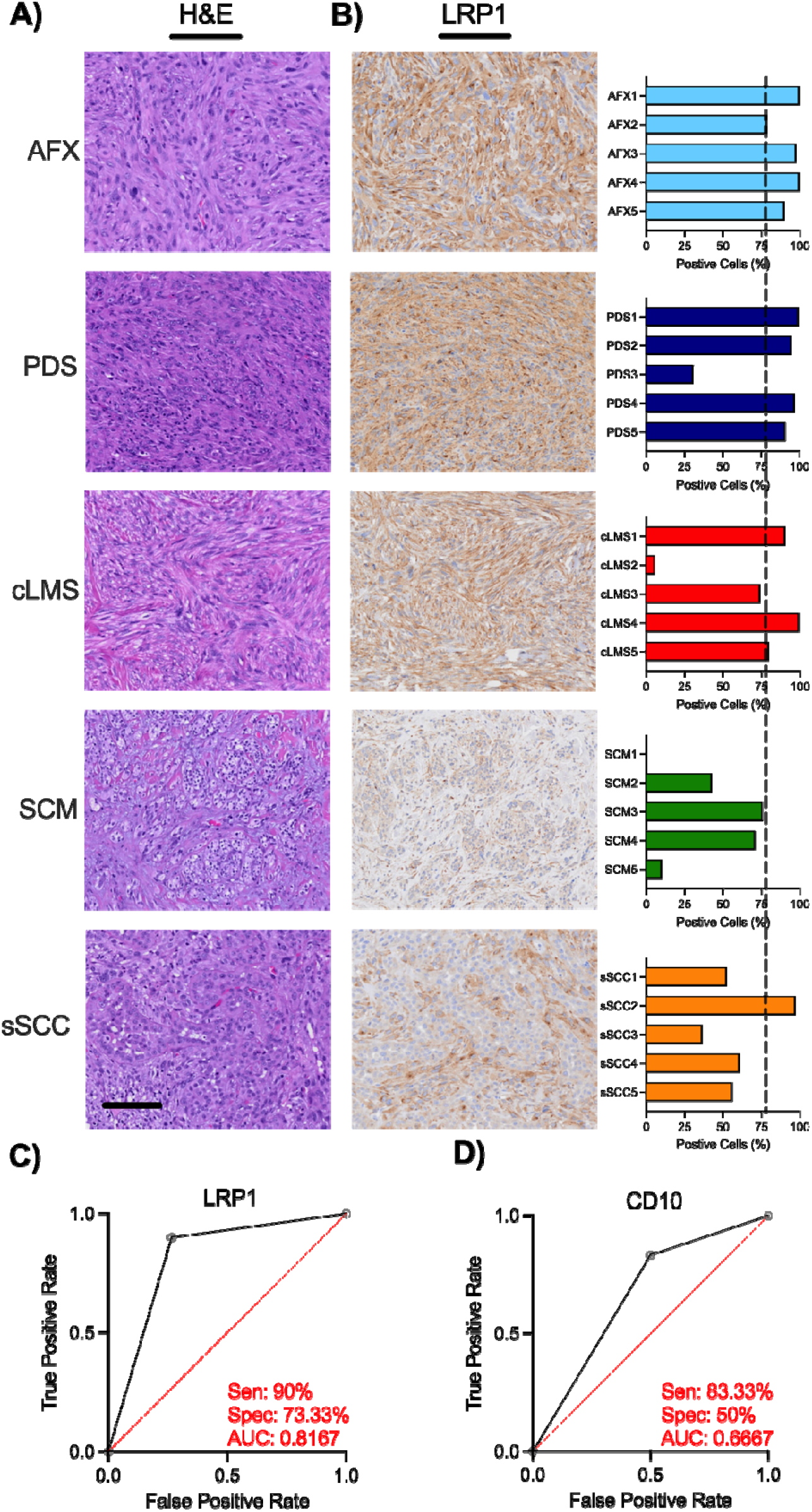
Immunohistochemical validation of LRP1 as a diagnostic biomarker for AFX/PDS. A) Representative brightfield images of H&E-stained tumor sections for each tumor type in the study: AFX, PDS, cLMS, SCM, and sSCC. Scale bar is 100 microns. B) Representative brightfield images of LRP1 immunohistochemistry in each tumor subtype with quantification of percent positive cells on the right. Dotted vertical line demarcates the threshold to be considered positive, at 77% positive cells. C) Reciever Operating Characteristic (ROC) analysis of C) LRP1 and D) CD10 (data extracted from Buonaccorsi *et al*. 2012) within the differential. Sensitivity (Sen), Specifity (Spec), and Area Under the Curve (AUC) are displayed.

The value of LRP1 as a sole diagnostic biomarker is limited by positive staining in cLMS (AFX/PDS vs cLMS AUC = 0.65; AFX/PDS vs SCM or sSCC AUC = 0.95 and 0.85, respectively) (Supplemental Figure 3A); however, it may be clinically useful if used in conjuction with markers to rule out cLMS, such as smooth muscle actin (SMA) or desmin (13). Of note, LMS was not included in the proteomic dataset that we used to filter candidate markers. Further studies in cLMS will improve our ability to distringuish this differential. Furthermore, use of LRP1 and LTBP2 in concert improves specificity from 73% to 93% (Supplemental Figure 3B-C). While these candidate biomarker have individual limitations, the interpretation of LRP1 expression in conjunction with other clinical tests like LTBP2, SMA, and desmin can improve our ability to resolve the tumors within this differential. Additionally, this study highlights the value of analyzing scRNA-seq and proteomics to identify biomarkers in rare tumors.

## Data Availability

All sequencing and proteomic data are publicly available. Accession numbers are available in the Supplemental Methods.

## Acknowledgements

We would like to thank Laura Hoaglin, Nathan Goldstein, Dr. Richard Wang, and Dr. Yiqun Shellman at CU AMC, Drs. Gary Hon/lab, Doug Strand, and Aya Alame at UTSW.

## Funding

This work was supported by NIH Training Grant 5T32AR007411-40 and the CU Cancer Center Support Grant (P30CA046934).

## Contributions

JJL and JCK designed this experiment. JCK, JJL, MMB, GTS performed data analysis and generated figures. GAH and AB performed case identification and confirmation of pathology. JJL and JCK drafted the manuscript. All authors reviewed and revised the manuscript.

## Notes

### Competing Interest Statement

The authors have declared no competing interest.

