## Supplemental Methods for "Proteomics and single-cell transcriptomics identify LRP1 as a diagnostic biomarker for atypical fibroxanthoma and pleomorphic dermal sarcoma"

**Supplementary Methods:**

**Single-cell RNA sequencing:**

Publicly available datasets for cSCC (GSM5788498, GSM5788500, GSM5788502), melanoma (GSM6622299, GSM6622300, GSM6622301), and uterine LMS (GSM7634954) were utilized in conjunction with single-cell RNA sequencing from 3 AFX and 2 PDS by our group (GSE281096 and GSE281529). Single-cell data sets were filtered for cells expressing >800 genes, genes that were expressed in > 20 cells, and mitochondrial reads. Samples were all normalized for read count and expression values were log transformed. Batch correction was performed using Scanorama. After batch correction, datasets were concatenated prior to principal component analysis and UMAP generation. Tumor populations from each dataset were called and genes were ranked by expression through logistic regression. The top 100 genes enriched in AFX/PDS were used as input for Gene Ontology (GO) analysis. The top 100 genes were then filtered for > 7-fold enrichment over other tumor types and minimal RNA expression in normal skin (GSE86571), which resulted in 12 candidate biomarkers for downstream analysis.

**Analysis of bulk proteomic data:**

Bulk proteomic data for PDS, SCC, and Melanoma were acquired from Klein *et al.,* 2024 (PMID 38822058). 6451 proteins from the Klein dataset were upregulated in PDS over SCC and melanoma. A One-way ANOVA was performed to determine statistically upregulated proteins in PDS. Out of the 6451 proteins reported to be upregulated in PDS, 3099 surpassed the threshold for statistical significance after correcting for multiple comparisons. The 3099 proteins statistically upregulated in PDS were ranked and used as input for Gene Ontology analysis.

**Immunohistochemistry**

A total of 25 formalin-fixed paraffin-embedded tumors (n = 5 per sarcoma subtype) were cut into 5 μM sections and placed onto positively charged glass slides by ProPath, a national pathology group. Slides were dewaxed using xylene and rehydrated with ethanol prior to a quenching endogenous peroxidase using 0.3% hydrogen peroxide for 20 minutes. Antigen retrieval was performed using Citrate-based Antigen Unmasking Solution (Vector Labs #H3300) for 20 minutes at medium lower power in the microwave. Tissue sections were blocked using 2.5% normal horse serum (Vector Labs #MP7401) for 40 minutes before an overnight incubation in rabbit anti-LRP1 (HPA022903, 1:250 diluted in 2.5% normal horse serum) or rabbit-LTBP2 (ThermoFisher #PA5-51930, 1:500 diluted in 2.5% normal horse serum) at 4C. The following day, tissue sections were washed twice with PBS prior to a 30-minute incubation with ImmPRESS HRP Horse Anti-Rabbit IgG (Vector Labs #MP7401). Slides were washed twice in 1x PBS before chromogen development using ImmPACT DAB Substrate Kit (Vector Labs #SK4105). Nuclei were counterstained using hematoxylin.

**Imaging and quantification of biomarker expression**

Brightfield images of stained tissue sections were obtained at 20x magnification on the Olympus VS200 slide scanner using Olympus OlyVIA v4.1 software for acquisition. Tumor area was annotated prior to quantification. Percent positive cells were determined using QuPath 0.6.0 and used to generate receiver operating characteristic (ROC) curves. Threshold for percent positive cells was set per IHC stain to maximize accuracy.

**Statistics**

Statistical analyses were performed using GraphPad Prism10, R studio, or Python. Comparisons between more than two groups were analyzed using a One-way ANOVA corrected for repeated measures. *p* < 0.05 was considered to mark statistically significant difference amongst cohorts.
