## Supplemental Figures for "Proteomics and single-cell transcriptomics identify LRP1 as a diagnostic biomarker for atypical fibroxanthoma and pleomorphic dermal sarcoma"

**Supplemental Figures:**
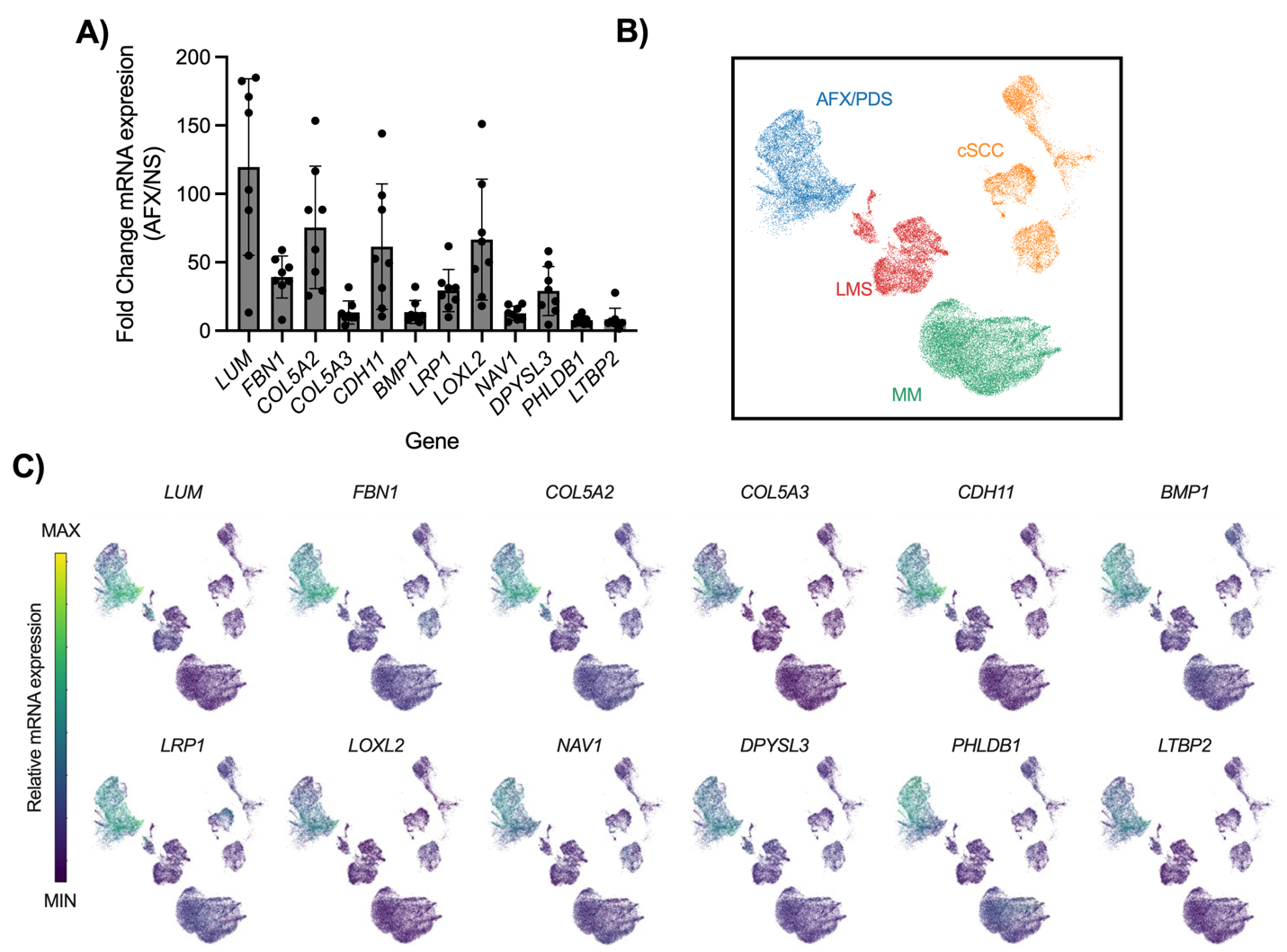

**Supplemental Figure 1: Analysis of candidate biomarker gene expression in normal skin and AFX/PDS cells.** A) The fold change in mRNA expression of each candidate gene in AFX over normal skin (NS) (Extracted from Lai *et al.)*. B) Dimensional reduction and visualization of scRNA-seq extracted from Klein *et al.* with AFX/PDS in blue, cSCC in orange, LMS in red, and MM in green. C) Feature plots of candidate biomarker expression overlayed onto the UMAP from Figure 1B.

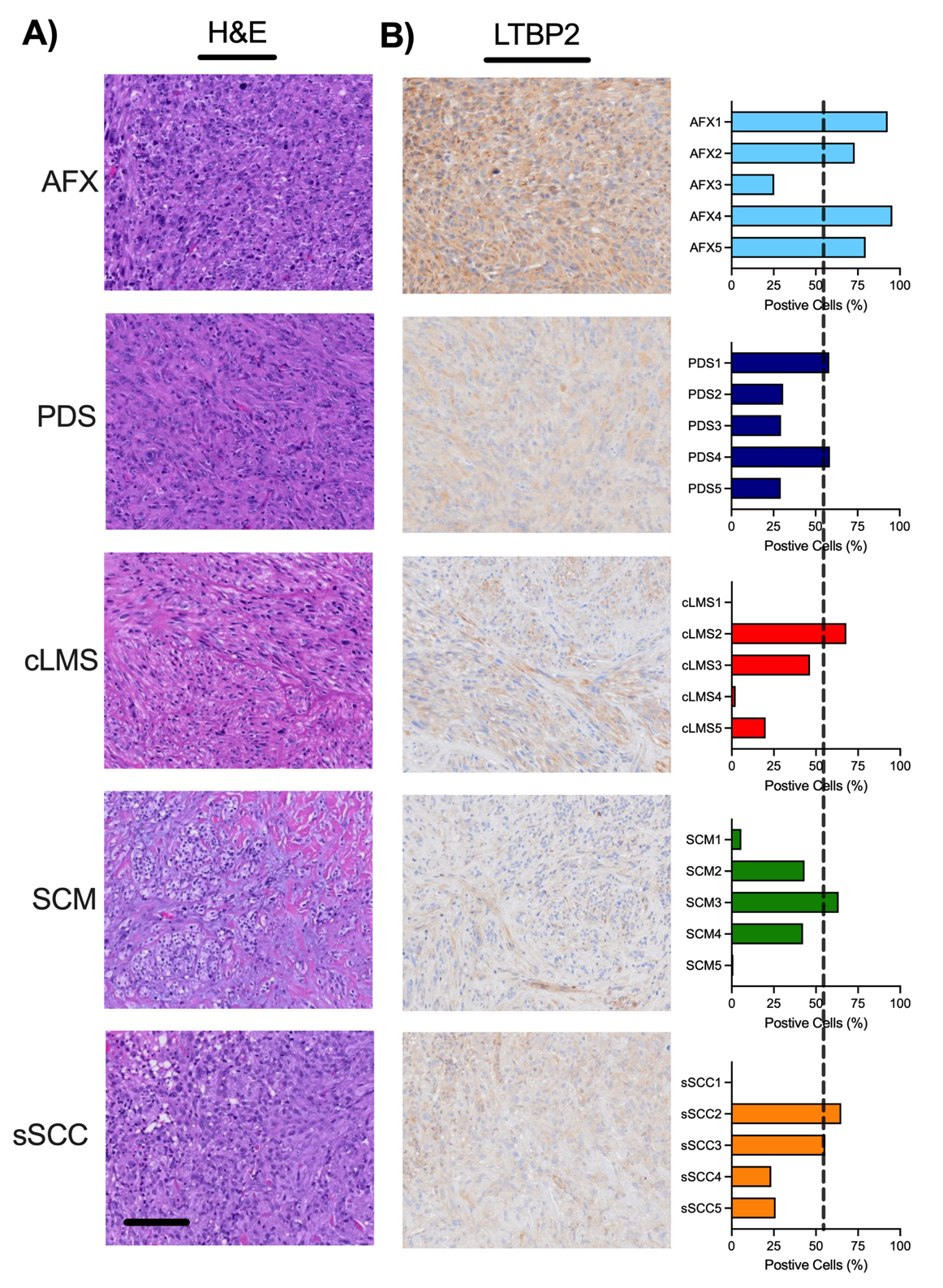

**Supplemental Figure 2: Immunohistochemical validation of LTBP2 as a diagnostic biomarker for AFX/PDS.** A) Representative brightfield images of H&E-stained tumor sections for each tumor type in the study: AFX, PDS, cLMS, SCM, and sSCC. Scale bar is 100 microns. B) Representative brightfield images of LTBP2 IHC in each tumor subtype with quantification of percent positive cells on the right. Dotted vertical line demarcates the threshold to be considered positive, at 57% positive cells.

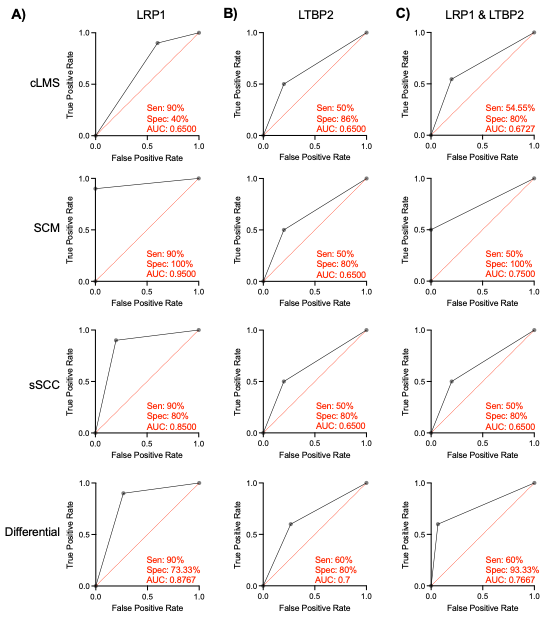

**Supplemental Figure 3: Receiver Operating Characteristic (ROC) analysis of LRP1, LTBP2, and the combination by tumor type.** ROC curves are displayed comparing AFX/PDS to each individual tumor type and the complete differential for A) LRP1, B) LTBP2, and C) LRP1 & LTBP2. Sensitivity (Sen), Specificity (Spec), and Area under the curve (AUC) are displayed for each curve.
